# Optical activation of midbrain dopamine neurons: do high and low stimulation frequencies produce functionally different effects?

**DOI:** 10.1101/2025.04.09.643378

**Authors:** Jacques Voisard, Marie-Pierre Cossette, Andreas Arvanitogiannis, Peter Shizgal

## Abstract

Millard et al. propose^1^ that 20- and 50-Hz stimulation of optically excitable, midbrain dopamine neurons triggers functionally distinct effects: The “physiological” firing elicited by a 20-Hz train mimics a reward-prediction error (RPE) whereas the “supra-physiological” firing induced by a 50-Hz train creates “a sensory event that acts as a reward in its own right.” Only the 50-Hz trains supported vigorous responding in the face of elevated response costs and sufficed to produce reward-specific Pavlovian-Instrumental Transfer (PIT). Nonetheless, the 20-Hz trains did produce unblocking. Here, we propose a more parsimonious account of these findings that is consistent with the results of prior experiments on operant responding for rewarding optical^2–4^ or electrical^5–9^ stimulation and with a new theory^10,11^ of the role of dopamine signaling in associative conditioning: 1) The effects of the 20- and 50-Hz trains differ quantitatively rather than qualitatively. They are mapped onto a single dimension, reward intensity, which reflects the aggregate release of dopamine in the critical terminal field(s). 2) The non-linear dependence of operant responding on both the aggregate dopamine release and the response cost^3^ can explain why responding for the 20- and 50 Hz trains diverged as response cost grew and predicts that this divergence can be eliminated by increasing the number of dopamine neurons stimulated. 3) Terminal-field dopamine concentration provides no information about pulse frequency per se. 4) Operant responding, PIT, and unblocking all depend on reward intensity, but the form and parameters of this dependence can differ, thus generating the observed contrasts in the impact of the 20- and 50-Hz trains. 5) The tested range of experimental conditions is too narrow to specify what constitutes “physiological” firing.

---

Psychophysical data, such as those in Figure 1^3^, imply that the rewarding effects of optogenetic stimulation of midbrain dopamine neurons are determined by aggregate dopamine release rather than by the pulse frequency per se. The data in Figure 1 were obtained from a TH-Cre rat expressing Channelrhodopsin-2 in midbrain dopamine neurons^3^; the rat pressed a lever to trigger delivery of 1 s pulse trains via an optical fiber aimed at the dopaminergic cell bodies. When the optical power was 10 mW, the rat worked at a mean rate of 23 responses s^-1^ (dotted line) for 50-Hz trains (dashed line), but when additional dopamine neurons were recruited by doubling the optical power, 18-Hz trains (dash-dot line) were able to produce the same level of behavior. Thus, a 20-Hz train can be as, or even more, effective than a 50 Hz train, provided that a sufficient number of neurons are recruited.

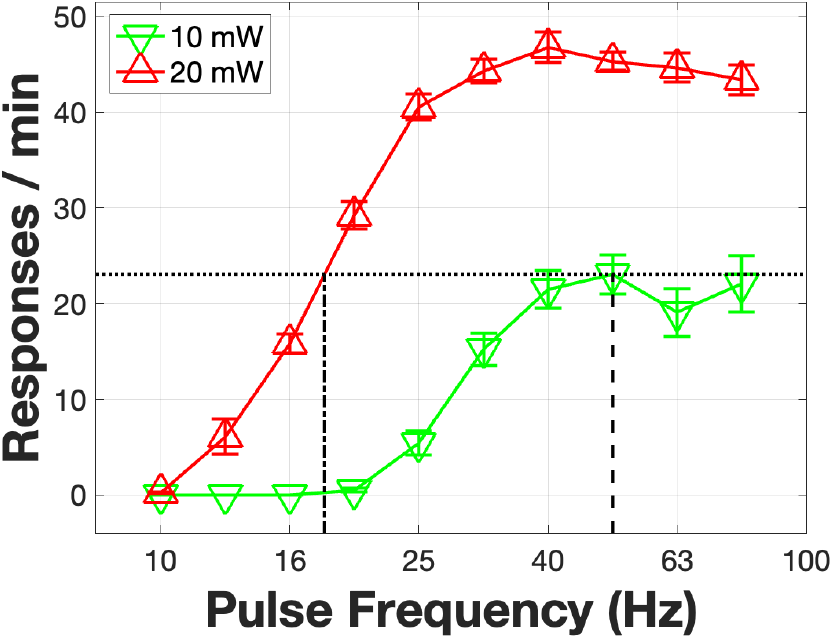

Millard et al. varied the response cost as well as the pulse frequency. We have done so previously in TH-Cre rats that earned optical stimulation of midbrain dopamine neurons^3^ by depressing a lever until the cumulative lever-depression time met a time-cost criterion (the “price”). Representative data from that study are shown in Figure S1. Reward seeking, measured as the proportion of trial time spent working for the stimulation (“time allocation”), varied steeply and non-linearly with both price and pulse frequency. The data are very well described by the fitted surface, which was generated by the “reward-mountain” model^3,12^. The surface in Figure S1 is reproduced in the left panel of Figure 2 along with the points on the surface corresponding to 20- and 50-Hz pulse frequencies delivered at low, intermediate, and high prices. These predicted values of time allocation are shown as inverted pink pyramids (20 Hz) and upright red pyramids (50 Hz), respectively. (Please see the extended methods in the supporting-information file for details.) The predicted time-allocation values in the left panel of Figure 2 reproduce the effect reported by Millard et al.: responding for the 20-Hz trains is robust at the lowest price, but falls off systematically as the price is increased, whereas responding for the 50-Hz train remains high at all three prices. Unlike the proposal of Millard et al., the mountain model entails no bifurcation of function on the basis of pulse frequency. Instead, aggregate excitation is mapped into reward intensity, which is scaled by reward cost and then translated into behavior by a variant of the single-operant matching law^3,13^.

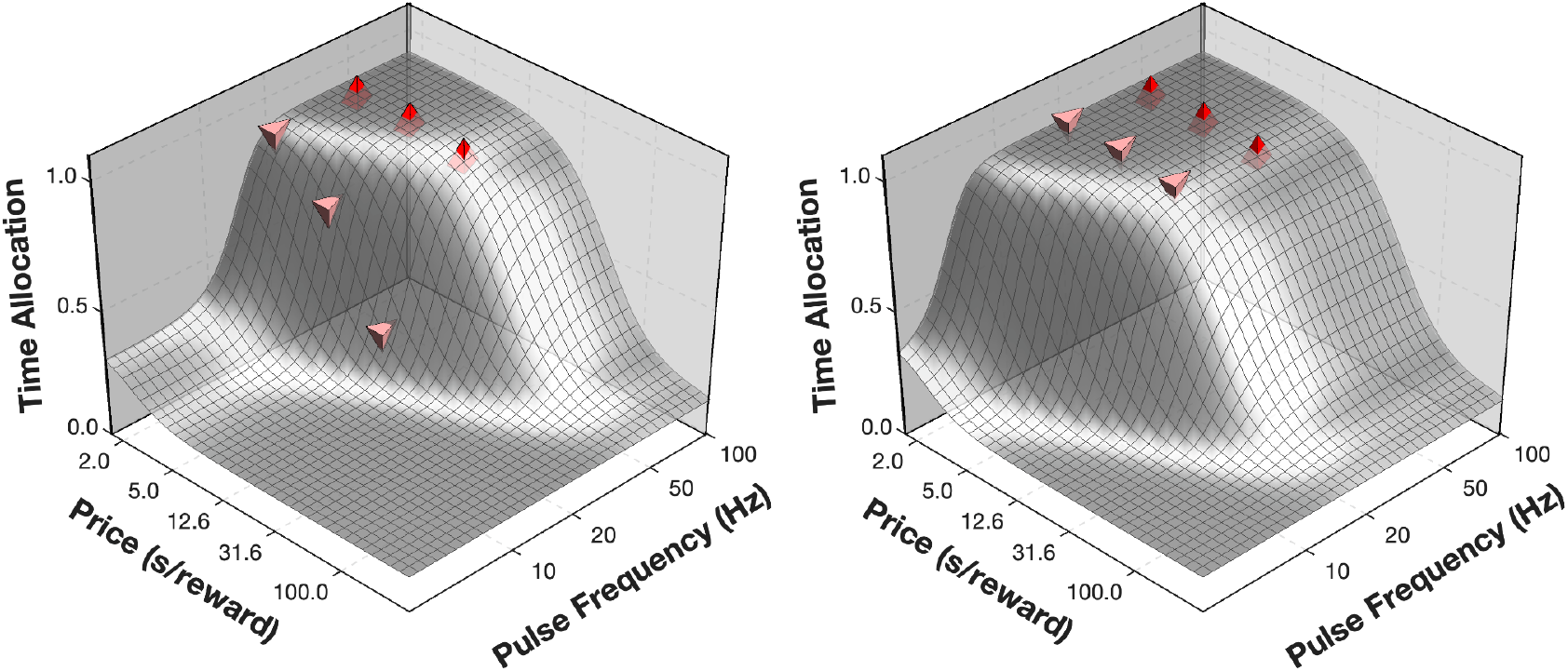

Increasing the size of the stimulated neuronal population shifts the reward mountain along the pulse-frequency axis towards the origin^13^ because lower pulse frequencies suffice to maintain a given level of aggregate dopamine release when more neurons contribute (Figure S2c). That effect is simulated in the right panel of Figure 2: The time-allocation values for the 20-Hz trains have now climbed onto the plateau of the shifted mountain, thus eliminating the differences in responding for the 20- and 50-Hz trains. In contrast, Millard et al. imply that responding for 20-Hz trains will necessarily be weak at elevated reward costs and less than responding for 50-Hz trains. (Please see the extended-methods section of the supporting-information file for an explanation of how the shifted surface and the data points were generated.)

Figure 2 illustrates how the choice of experimental parameters can dramatically alter the relationship between behavior, response cost, and pulse frequency. Analogously, the differential effectiveness of the 20- and 50-Hz trains in Millard et al.’s PIT test, and of the 20-Hz train in the PIT and unblocking tests, could stem from the particular stimulation parameters and task-variable settings employed, which can position the curves mapping pulse frequency into the various behavioral outcomes at different locations along the pulse-frequency axis.

For 20-Hz and 50-Hz trains to produce functionally different effects (e.g., reward versus RPE), information about the pulse frequency per se must be relayed to the synaptic targets of the dopamine neurons. However, fast-scan cyclic voltammetric (FSCV) recordings of dopamine release in the Nucleus Accumbens (NAc) argue that this information is ***not*** preserved in terminal-field dopamine concentrations (Figure S2, methodological details in the extended-methods section of the supporting-information file). These recordings show that optogenetically induced changes in terminal-field dopamine concentrations reflect both the number of activated neurons and the rate at which optical pulses are delivered. A given concentration change can be produced by altering the optical power and hence the number of activated neurons (Figure S2c), by altering the pulse frequency and hence the amount of dopamine released per neuron^4,14^ (Figure S2d), or both. Thus, the extracellular concentration of dopamine impinging on the postsynaptic receptors near these NAc FSCV electrodes reflected aggregate dopamine release and conveyed no information about pulse frequency per se.

We now address the issue of nomenclature. To forestall satiation, the rewards employed in electrophysiological recording studies of dopaminergic responses are typically small; ethical constraints require that food or water deprivation be mild. Such conditions can capture only a thin slice of the conditions and experiences encountered in the wild and do not provide an adequate basis for defining a boundary between “physiological” and “supra-physiological” firing. Thus, we dispute the category names adopted by Millard et al. and recommend that their use be suspended. That ∼50-Hz responses to air puffs have been recorded from putative dopamine neurons in monkeys^15^ provides further grounds for reconsidering the categorization and labeling.

Millard et al. dichotomize an RPE and a “reward in its own right” whereas a new theory^10,11^ treats the contributions of dopamine and natural rewards to the learning of causal relations as commensurable and additive. On that view, a common dopaminergic mechanism imbues reward-predicting stimuli with predictive value and retrodicts the actions that have led to reward delivery. The input dimension for the dopaminergic contribution to causal learning is the aggregate release of dopamine in the relevant terminal field(s).

## Acknowledgements

The authors thank Matt Gardner, Giovanni Hernandez, Mihaela Iordanova and Vijay M.K. Namboodiri for their helpful comments.

