## Supporting information for "Optical activation of midbrain dopamine neurons: do high and low stimulation frequencies produce functionally different effects?"

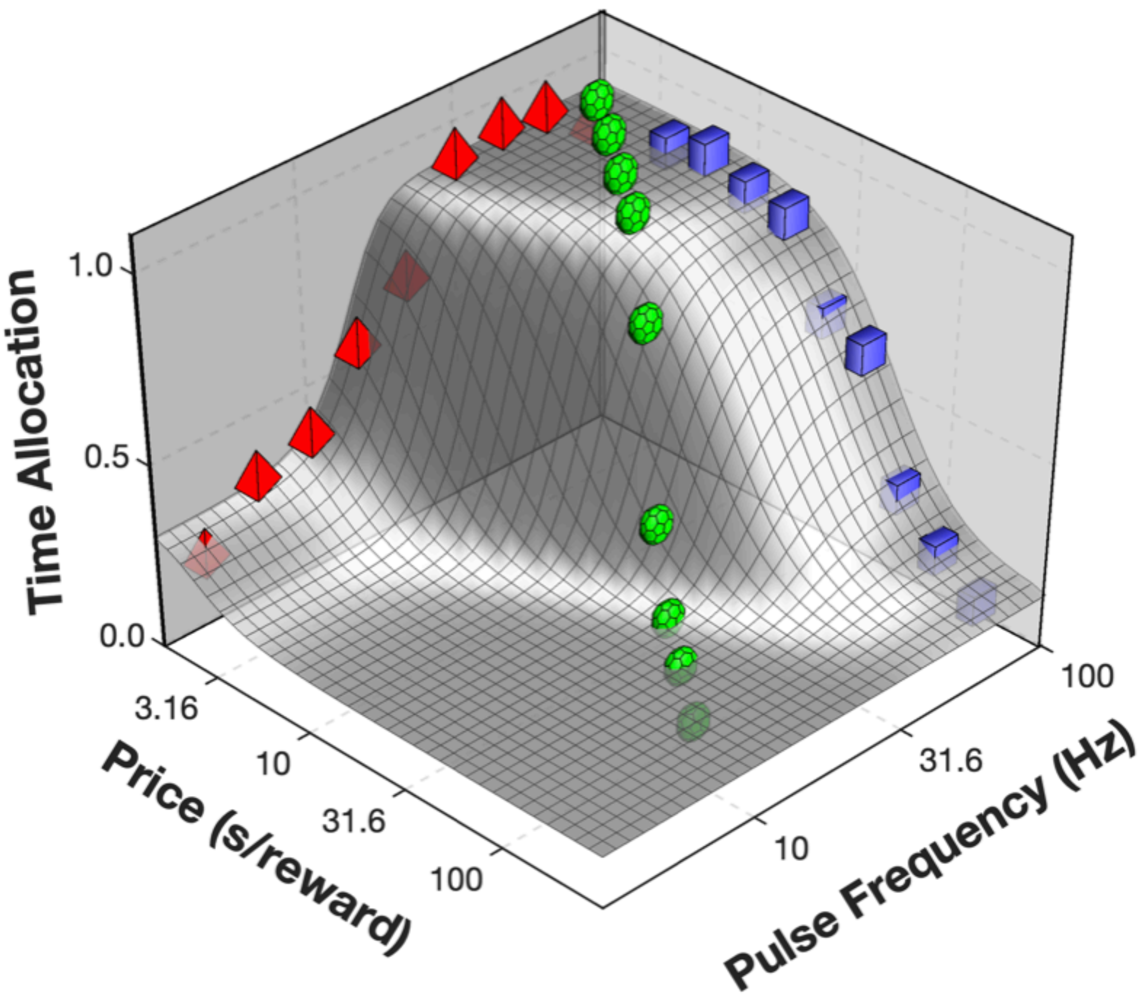

Time Allocation
1.0
0.5
0.0
3.16
10
31.6
100
Price (s/reward)
100
31.6
10
Pulse Frequency (Hz)


Figure S1

**Figure S1**: Previously published self-stimulation data from a rat (BeChR28) working for rewarding optical stimulation of midbrain dopamine neurons expressing ChR2[3]. The rat had been trained to hold down a lever to earn 1-s stimulation trains of 5-ms pulses, which were delivered once the cumulative hold-down time reached a criterial value (the “Price”). Thus, the X-axis represents the opportunity cost (“Price”) of the reward, the Y-axis represents the pulse frequency, and the Z-axis represents the proportion of trial time during which the lever was depressed (“Time Allocation”). The data fall along three trajectories. Red tetrahedrons represent average time allocation at various pulse frequencies while the price was fixed at 2 s. Blue, solid rectangles represent average time allocation at various prices while the pulse frequency was fixed to a very high value. Green soccer balls represent average time allocation at pairs of prices and pulse frequencies chosen to orient their trajectory diagonally between those denoted in red and blue. The black line is the time-allocation contour that falls midway between the summit and the floor. The curved surface, the corner of a plateau that we dub the “reward mountain,” is the 3D fit of a model[3,5,6,7] that relates time allocation to the strength of the rewarding effect and the opportunity cost of the stimulation. Please see the original paper[3] for details.

*Citations refer to the reference list for the main text.*

### Dopamine concentration as a function of pulse frequency and optical power

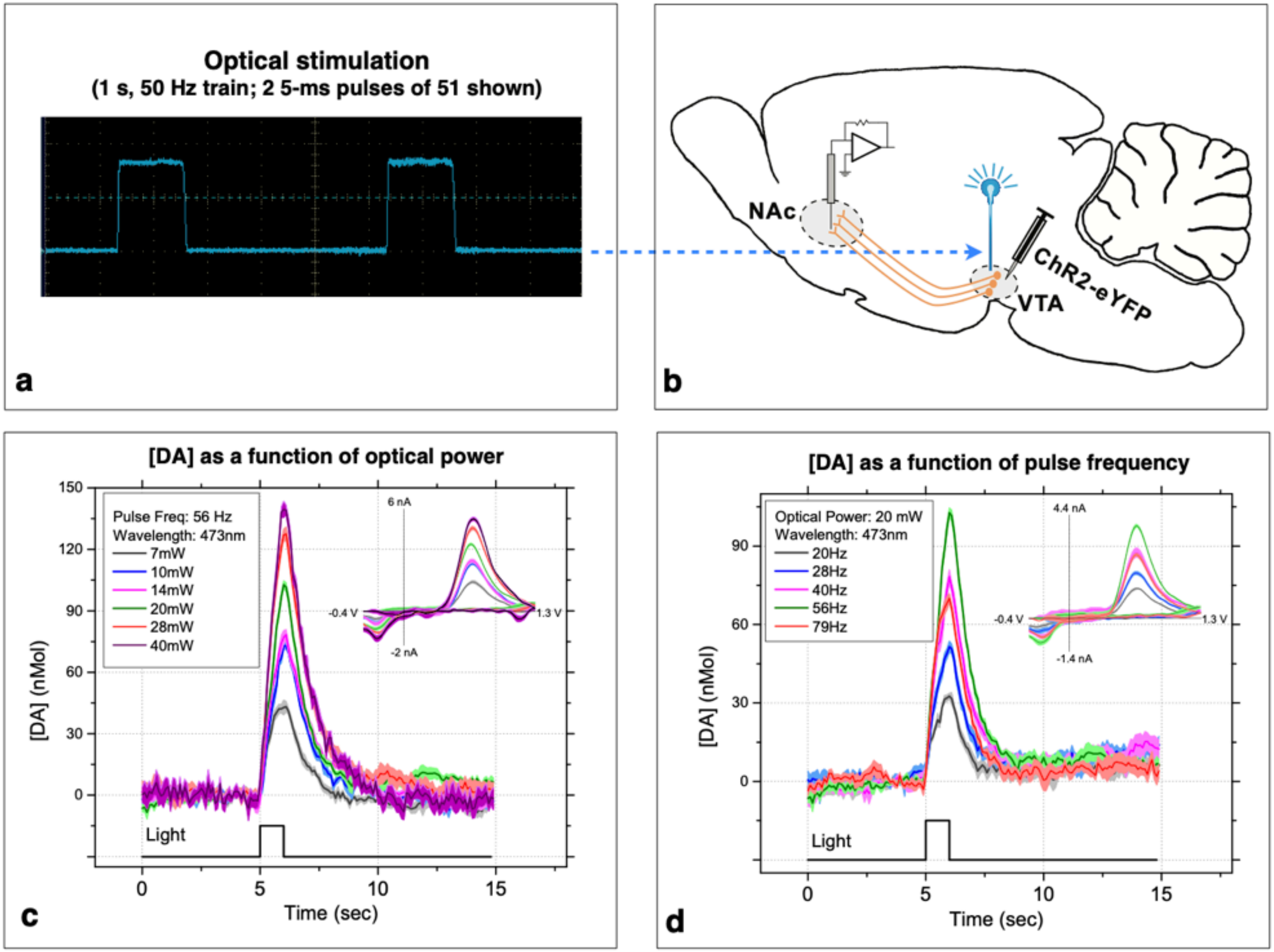


Figure S2

**Figure S2:** a) Sample output from the 473-nm DPSS laser used to drive the dopamine release described in panels c and d. The laser operated continuously, and its ouput was chopped by a mechanical shutter. See Extendend methods below for details.

b) Sagittal sketch of a rat brain showing the setup used to obtain the data in panels c and d. The optical fiber (cyan) was aimed at the Ventral Tegmental Area (VTA), and the carbon-fiber electrode was aimed at the Nucleus Accumbens (NAc). The VTA->NAc projections of Channelrhodopsin2-expressing dopamine neurons are represented in orange. See Extendend methods below for details.

c) FSCV recordings of NAc dopamine transients driven by optical stimulation of the VTA at different optical powers in a urethane-anesthetized TH-Cre rat. The inset in the upper right is a voltammogram showing the recorded oxidation and reduction currents. See Extendend methods below for details.

d) FSCV recordings of NAc dopamine transients driven by optical stimulation of the VTA at different pulse frequencies in a urethane-anesthetized TH-Cre rat. See Extendend methods below for details and Covey & Cheer (2019)[14] for analogous results.

It is evident from panels c) and d) that a transient of a given amplitude could be generated by multiple pairs of pulse frequencies and optical powers. A given change in the amplitude of the transient could be produced by altering the pulse frequency, the optical power, or both. Thus, the extracellular dopamine concentration does not provide information about either the pulse frequency per se or the optical power per se. Instead it reflects the combined effect of these two variables. The pulse frequency determines the impulse flow in the stimulated neurons, whereas the optical power determines the number of neurons that are fired. Thus, the pulse frequency and optical power jointly determine the aggregate release of dopamine in the region surrounding the FSCV electrode and the consequent changes in dopamine concentration that are sensed by the carbon-fiber electrode.

These are previously unpublished data that could not have been seen by Millard et al. prior to the preparation of their paper.

### Extended methods

**Figure 1**: This figure shows a subset of the data in a previously published figure[3] (Figure S2 in the supporting-information file for that paper, which describes results from rat Bechr19). For clarity, the curves in the original figure for the 2.5, 5, and 40 mW optical powers were removed, leaving only the curves obtained at 10 and 20 mW. The MATLAB (R2024b) code used to produce Figure 1 can be found in the Open Science Foundation project for this paper.

**Figure 2**: The surface in the left panel is taken from a previously published figure[3] (Figure S22 in the supporting-information file for that paper, which describes the results from rat Bechr28). That supporting-information file also includes a derivation of the equation that generated the surface that best fit the data from that rat (Equation S38) as well as the values of the fitted parameters (Table S15). The equation in question is the conditioned-reward variant of the reward-mountain model. That model translates the pulse frequency and time-cost ("Price") of the rewarding optical-stimulation train into the proportion of trial time that the rat worked for the stimulation ("Time Allocation").

The simulated data points in the left panel were positioned at the locations on the fitted surface corresponding to the following values of the two independent variables:

| Pulse Frequency | Low Price | Intermediate Price | High Price |
|---|---|---|---|
| **20 Hz** | 2 s | 5 s | 12.6 s |
| **50 Hz** | 2 s | 5 s | 12.6 s |

Positioning the data points at these locations is equivalent to solving the conditioned-reward variant of the reward-mountain equation, using the independent-variable values in the table and the fitted parameter values for rat Bechr28.

The surface in the right panel of Figure 2 was obtained by shifting the surface in the left panel along the pulse-frequency axis by 0.386 common-logarithmic units. That shift is what the mountain model predicts when an increase in optical power and/or pulse duration boosts the number of stimulated dopamine neurons by a factor of 2.432 ($10^{0.386}$).

The parameter that positions the surface along the pulse-freqency axis is called $F_{Pulse_{hm}}$; it is the pulse frequency at which the intensity of the stimulation-induced rewarding effect is half-maximal[3]. The shift that produced the surface in the right panel of Figure 2 reduces the fitted common-logarithmic value of $F_{Pulse_{hm}}$ for rat Bechr28 by 0.386 (equivalent to dividing $F_{Pulse_{hm}}$ by 2.432). The simulated data points in the right panel of Figure 2 were positioned at the locations on the shifted surface corresponding to the pulse frequencies and prices in the above table.

The MATLAB (R2024b) code used to produce Figure 2 can be found in the Open Science Foundation project for this paper.

**Figure S2** shows data from an acute Fast-Scan Cyclic Voltammetry (FSCV) recording session carried out in an anesthetized TH-Cre rat in which mid-brain dopamine neurons were activated optogenetically. The following methods were used to prepare the subject and gather the data.

The subject was a TH-Cre, heterozygous, male, Long-Evans rat descended from founders provided by Karl Deisseroth. An initial surgery was performed to express the excitatory opsin, Channelrhodopsin2 (ChR2), in midbrain dopamine neurons. The rat was anesthetized with a mixture of ketamine hydrochloride (87 mg/kg) and xylazine hydrochloride (13 mg/kg). An s.c. injection of atropine sulphate was given to reduce bronchial secretions (0.05 mg/kg), an s.c. injection of buprenorphine to alleviate pain (0.05 mg/kg), and an s.c. injection of penicillin procaine to prevent infection (0.3 cc/rat). ‘HypoTears’ (Polyvinyl alcohol 1% [10 mg/1 mL]) was applied to the cornea for protection. Once deep anesthesia had been achieved, the fur over the skull was shaved, the exposed skin was disinfected and treated topically with a local anesthetic (50:50 v/v mixture of 2% lidocaine and 0.5% bupivicaine). After an anaesthetic ointment (Xylocaine) was applied to the external auditory meatus, the rat was mounted in a stereotaxic frame (Kopf Instrument, Tujunga, CA), and its snout was placed in a nose cone so that volatile anesthetic could be administered continuously throughout the surgery (0.5–1.5% isoflurane). Body temperature was controlled throughout the surgical procedures with a heating pad. An s.c. injection of Ringer’s solution (6 mL/kg) was given every hour. An incision was made in the scalp over the midline, the wound margins were retracted, the skull was levelled, and the periosteum above the ventral tegmental area (VTA) was removed. Burr holes were drilled above the VTA to provide access for the 28-gauge injector that was to be used for viral transfection of the VTA dopamine neurons. The injector was connected to a Hamilton syringe, driven by a Harvard pump.

The viral vector was a serotype-5 adeno-associated virus (AAV5) containing an EF1a-DIO-hChR2(H134R)-EYFP-WPRE transcript suspended in phosphate buffered saline. Transfection was initiated by injecting 0.5 $\mu$l of the vector-containing solution at each of six locations above and within the VTA at a rate of 0.1 $\mu$L min$^{-1}$, with a pause of 10 min separating each injection. The coordinates for the injections were: {AP: 5.4, ML: 1.0, DV: 8.3, 7.7, 7.2}; {AP: 6.2, ML: 1.0, DV: 8.3, 7.7, 7.2}. Following completion of the injections, the injector was slowly withdrawn, the burr holes sealed with sterile bone wax, the scalp incision was sutured, the rat was removed from the stereotaxic and then observed until normal locomotion was resumed. At least four weeks were provided for ChR2 expression and transport. (We are unable to locate the records specifying the exact duration of this interval.)

The carbon-fiber working electrode was constructed by threading a 7 $\mu$m diameter carbon fiber (Thorne, Amoco Corporation, Greenville, SC, USA) through a single-barrel, borosilicate glass capillary ({ID:0.40 mm; OD: 0.60 mm};. A-M Systems, Carlsborg, WA, USA). The bare carbon fiber protruded from the end of the silica tubing by 150-200 $\mu$m. The carbon fiber was sealed within the glass capillary by heating the capillary with a pipette puller (PUL-1, WPI, Sarasota, FL, USA). A wire covered with silver paint (GCElectronics, Rockford, IL, USA) was inserted in the capillary to make contact with the carbon fiber and secured with shrink tubing coated with epoxy. The free end of the wire was terminated in a gold-plated Amphenol connector.

The implant to be used to activate the ChR2-expressing dopamine neurons was fashioned from a step-index optical fiber (300 ± 6 μm core, 325 ± 10 μm cladding, NA = 0.39). One end of the fiber was terminated in a 2.5-mm steel ferrule, glued in place and polished.

The carbon-fiber working electrode was pre-calibrated in an electrochemical cell connected to a 6-port pressure injection valve (Upchurch Scientific, Oak Harbor, WA). The electrode was cleaned prior to calibration with 2-propanol containing activated carbon. Artificial cerebrospinal fluid (aCSF; 145 mM Na+ , 2.7 mM K+ , 1.22 mM Ca2+ , 1.0 mM Mg2+ , 150mM Cl–, 0.2mM ascorbate, 2mM Na2HPO4, pH 7.4±.05) was passed through the electrochemical cell using a pump (Multi-Phaser, YA-12, Yale Apparatus, Holliston, MA). The loop injector was loaded with DA standards at concentrations of 100nM, 200nM, 500nM, and 1000nM. DA was dissolved in ASCF. Only electrodes with linear responses to standards of increasing concentration were kept for the in vivo voltammetric measurements.

To prepare for the FSCV recording session, the rat was anesthetized with urethane (1.5 g/kg) and mounted again in the stereotaxic frame. The scalp incision was opened, the skin was retracted, and the periosteum over the NAc was removed. A burr hole for the carbon-fiber working electrode was drilled above the NAc (AP: 1.7 mm; ML: 1.0 mm) and the dura mater below it was opened. To accommodate the sintered Ag/AgCl reference electrode (In Vivo Metrics, Healdsburg, CA, USA), a second burr hole was drilled in the skull over a location (~AP: –7.0 mm; ~ML: 4.0 mm) contralateral to the burr hole over the NAc. The bone wax in the previously drilled burr holes over the VTA was removed, the skull between the two holes was drilled away, and the optical implant was lowered to the following coordinates: {AP: 5.8; ML: 1.0 mm; DV = 7.7 mm}. The reference electrode was then inserted so that at least a 4 mm length was within the hemisphere contralateral to the working electrode. A small bead of dental acrylic was used to fix the reference electrode to the skull. The working electrode was then lowered slowly to its starting position, 7 mm below the skull surface, and the reference electrode was inserted so that at least a 4 mm length was within the hemisphere contralateral to the FSCV electrode.

Background-subtracted cyclic voltammograms were generated at 10 Hz by applying a 8.5 ms triangular waveform that ramped from -0.4 V to +1.3V and back to -0.4 V at a scan rate of 400 V/s. The potential was held at -0.4 V between each scan to promote cation absorption at the surface of the carbon fiber. The waveform was generated using LabVIEW (National Instruments, Austin, TX) and a multifunction data acquisition board (PCI-6052E, National Instruments, Austin, TX). A PCI-6711E board (National Instruments, Austin, TX) was used to perform waveform acquisition and data collection. All potentials were measured with respect to the Ag/AgCl reference electrode using a headstage based on a design by Scott Ng-Evans (University of Washington). The output of the headstage was connected to an analogue input of the PCI-6052E board to monitor the current passing through the FSCV electrode. The FSCV data were logged by software written by M. L. A. V. Heien and R. M. Wightman (Department of Chemistry, University of North Carolina, Chapel Hill, NC, USA). A synchronization signal from the PCI-6711E board was sent to the external input of a multi-channel pulse generator (Master-8, A.M.P.I, Israel) and used to trigger 1-s trains of optical stimulation 5 s after the start of each recording.

The depth of the working electrode was adjusted until a) voltammograms were obtained with a reliable cyclic-voltammetry signature characteristic of DA, with a peak oxidation at roughly 0.6 V and a reduction peak at -0.2 V (see insets in Figures S2c and S2d) and b) further lowering failed to increase the amplitude of the evoked dopamine transient. The working electrode was then conditioned at 60 Hz for 10 min and allowed to stabilize at 10 Hz for 10 min. Six recordings, 15 s in duration, were obtained using each pair of optical-power and pulse-frequency values.

Peak detection to quantify DA transients was performed using a customized routine in MATLAB (The Mathworks, Natick, MA). The maximal change in peak oxidation currents was measured during the rising phase of the triangle wave. For conversion into molar concentrations, these peak currents were compared to post-calibration measurements carried out *in-vitro*. This conversion generates six vectors consisting of 150 peak-concentration estimates (10 Hz $\times$ 15 s, replicated 6 times) for each pair of optical-power and pulse-frequency values. The mean of standard error at each time point along these vectors were calculated and are shown in Figures S2c and S2d.
